# Quantifying small GTPase activation status using a novel fluorescence HPLC-based assay

**DOI:** 10.1101/2024.10.02.615746

**Authors:** Makoto Araki, Yukika Kasuya, Kaho Yoshimoto, Toshiaki Katada, Kenji Kontani

**Affiliations:** Department of Biochemistry, Meiji Pharmaceutical University, Tokyo 204-8588, Japan; Molecular Cell Biology Laboratory, Research Institute of Pharmaceutical Sciences, Faculty of Pharmacy, Musashino University, Tokyo 202-8585, Japan

**Keywords:** HPLC, small GTPase, RHEB, KRAS, sotorasib

## Abstract

Small GTPases play crucial roles in cellular signaling pathways, with their activation states tightly regulated between GDP-bound inactive and GTP-bound active forms. Dysregulation of these nucleotide-binding states, such as in oncogenic RAS, is implicated in diseases like cancer. Accurately quantifying these states in cells is thus crucial for deciphering their functional roles and regulatory mechanisms. However, current methods do not fully meet the necessary sensitivity and versatility, limiting their effectiveness in small GTPase analysis. Here, we present a highly sensitive HPLC-based assay with fluorescence detection (Fluor-HPLC), enabling precise quantification of guanine nucleotide-binding states in small GTPases. Applying this technique, we successfully quantified the guanine nucleotide-binding states of RHEB and KRAS at their endogenous expression levels. We demonstrated the utility of Fluor-HPLC by elucidating RHEB activation dynamics in response to insulin stimulation and amino acid availability. Furthermore, integration of Fluor-HPLC with syngeneic mouse models provided insights into KRAS activation dynamics in tumor tissues and evaluated the effectiveness of targeted therapeutics. Overall, this versatile method paves the way for investigating activation states and regulatory mechanisms of various small GTPases, potentially accelerating our understanding of their roles in cellular processes and disease pathogenesis.

## Introduction

Small GTPases act as molecular switches, modulating various intracellular signaling pathways through conformational changes between their inactive GDP-bound and active GTP-bound forms (Bourne et al., 1991; Vetter and Wittinghofer, 2001). In their GTP-bound state, small GTPases interact with effector proteins to regulate key biological processes such as cell proliferation, cytoskeletal organization, and intracellular trafficking. The guanine-nucleotide binding state of small GTPases is governed by the balance between the GDP-GTP exchange and GTP hydrolysis reactions, which are facilitated by guanine nucleotide exchange factors (GEFs) and GTPase-activating proteins (GAPs), respectively (Bos et al., 2007; Cherfils and Zeghouf, 2013; Vigil et al., 2010). As observed with the proto-oncogene product RAS, dysregulation of the guanine-nucleotide binding state is implicated in the pathogenesis of various diseases, including cancer (Hobbs et al., 2016; Teng, 2022). Therefore, elucidating the intracellular binding states of small GTPases is crucial for understanding their functions and roles in disease.

Numerous methods have been developed to analyze the guanine-nucleotide binding state of small GTPases in cells. However, these methods present challenges in terms of simplicity and versatility. For instance, metabolic labeling using [^32^P]orthophosphate can detect GDP and GTP with high sensitivity but requires radioisotope facilities and poses radiation risks to cells (Bond et al., 1999; Inoki et al., 2003; Kontani et al., 2002; Satoh et al., 1988). The pull-down assay, which employs effector proteins fused with GST or antibodies selectively recognizing the active form of small GTPases, is straightforward and convenient for investigating the activation status of small GTPases (Benard and Bokoch, 2002; Taylor et al., 2001). However, this approach requires well-characterized effector proteins and specific antibodies. FRET probes enable spatiotemporal analysis of the activation state of intracellular small GTPases, but developing specific FRET probes is a demanding and intricate process (Kim et al., 2019; Kiyokawa et al., 2011). Due to these limitations, only a limited number of small GTPases have had their intracellular activation states and regulatory processes adequately investigated. We have previously reported a method for analyzing the activation state of small GTPases by quantifying their bound guanine nucleotides using ion-pair reversed-phase high-performance liquid chromatography (IP-RP-HPLC) (Araki et al., 2021; Araki and Kontani, 2024). This approach allowed the analysis of guanine nucleotides bound to small GTPases immunopurified from cell lysates. However, ultraviolet spectrophotometric detection in HPLC required overexpression of the target small GTPases in cells to achieve detectable levels of bound guanine nucleotides.

In this study, we introduce a highly sensitive HPLC-based assay capable of analyzing the guanine-nucleotide binding states of various small GTPases. This method enables the assessment of RHEB and KRAS activation status at their endogenous expression levels and allows for the investigation of their responses to insulin stimulation and KRAS/G12C inhibitors. Additionally, we demonstrate that this method, when combined with syngeneic mouse tumor models, provides valuable insights into the guanine-nucleotide binding state of KRAS in tumor tissues and serves as an effective tool for evaluating the efficacy of KRAS/G12C inhibitors *in vivo*.

## Results

### Highly Sensitive Quantification of GDP and GTP Using Reversed-Phase High-Performance Liquid Chromatography with Fluorescence Detection (Fluor-HPLC)

We aimed to establish a highly sensitive HPLC-based assay for evaluating the guanine-nucleotide binding status of small GTPases at their native expression levels. We employed fluorescent derivatization of guanine nucleotides to achieve this, enabling their high-sensitivity analysis via an HPLC system. The derivatization involved 3,4-dimethoxyphenylglyoxal (DMPG), a compound known to react with guanine, its nucleosides, and nucleotides, as previously described (Ohba et al., 1994). We optimized the conditions of reversed-phase high-performance liquid chromatography with fluorescence detection (Fluor-HPLC) to separate and detect fluorescently derivatized GDP, GTP, and Ganciclovir (an internal standard for the fluorescence derivatization reaction), achieving retention times of 1.9, 2.5, and 6.9 minutes, respectively (Fig. 1 and *SI Appendix*, Fig. S1). Under these optimized conditions, the limit of quantification (LoQ) for GDP and GTP was approximately 2.2 fmol and 0.7 fmol, respectively, representing an approximately 100-fold improvement over our previous IP-RP-HPLC method (Table 1). Moreover, the correlation coefficient (*r*²) obtained from linear regression analysis within the calibration concentration range (2 fmol – 75 pmol) was ≥0.999 for both GDP and GTP.

**Figure 1.**
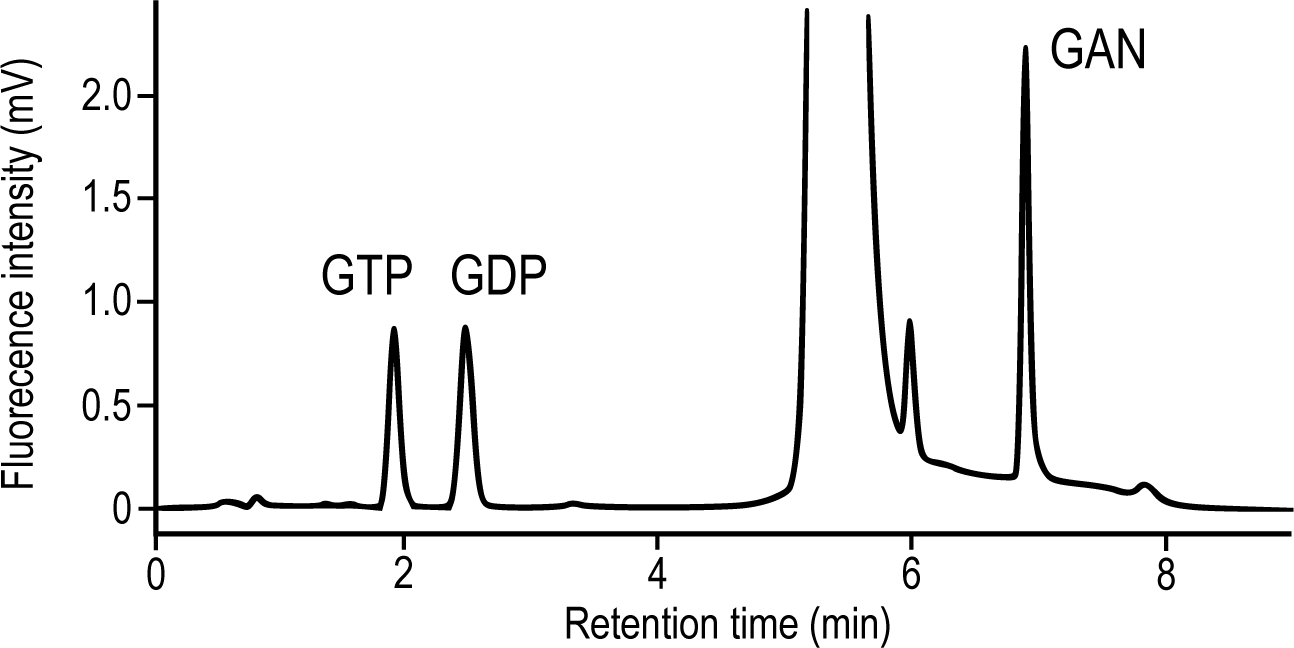
Fluor-HPLC separation of guanine nucleotides and ganciclovir. Representative chromatogram of a standard mixture of guanine nucleotides (GTP and GDP, each 200 fmol) and ganciclovir (GAN, 375 fmol, an internal control) by Fluor-HPLC. The compounds were detected using a fluorescence detector at an excitation wavelength of 400 nm and an emission wavelength of 510 nm. Peaks of GTP, GDP, and GAN were detected at 1.9, 2.5, and 6.9 min, respectively, on the chromatograms.

**Table 1.** Analytical data for the GTP and GDP standards using Fluor-HPLC.

|  | Retention time | CV<sup>a</sup> | LoQ<sup>b</sup> |
| --- | --- | --- | --- |
| GTP | 1.9 min | 4.1% | 2.2 fmol |
| GDP | 2.5 min | 1.4% | 0.7 fmol |

<sup>a</sup> coefficient of variation<sup>b</sup> limit of quantification

### Analysis of the Activation State of RHEB at Endogenous Expression Levels Using Fluor-HPLC

Next, we assessed the feasibility of analyzing guanine nucleotides bound to small GTPases using Fluor-HPLC. We focused on RHEB, a critical small GTPase that serves as a molecular switch controlling cellular responses to growth factors, nutrient availability, and stress stimuli (Inoki et al., 2003; Long et al., 2005; Saucedo et al., 2003; Stocker et al., 2003). FLAG-tagged RHEB was expressed in HeLa cells in a doxycycline (Dox)-dependent manner to ensure expression levels akin to endogenous RHEB (Fig. 2*A*). Following immunoprecipitation of FLAG-RHEB from cell lysates, guanine nucleotides bound to FLAG-RHEB were dissociated from proteins via heat denaturation and subjected to Fluor-HPLC analysis. Our results consistently enabled the quantification of GDP and GTP bound to FLAG-RHEB (Fig. 2*B*). Notably, the proportion of GTP-bound FLAG-RHEB was approximately 30%, aligning closely with findings reported for endogenous RHEB in a prior study (Im et al., 2002). These data indicate the utility of Fluor-HPLC in elucidating the activation state of RHEB at its endogenous expression levels.

**Figure 2.**
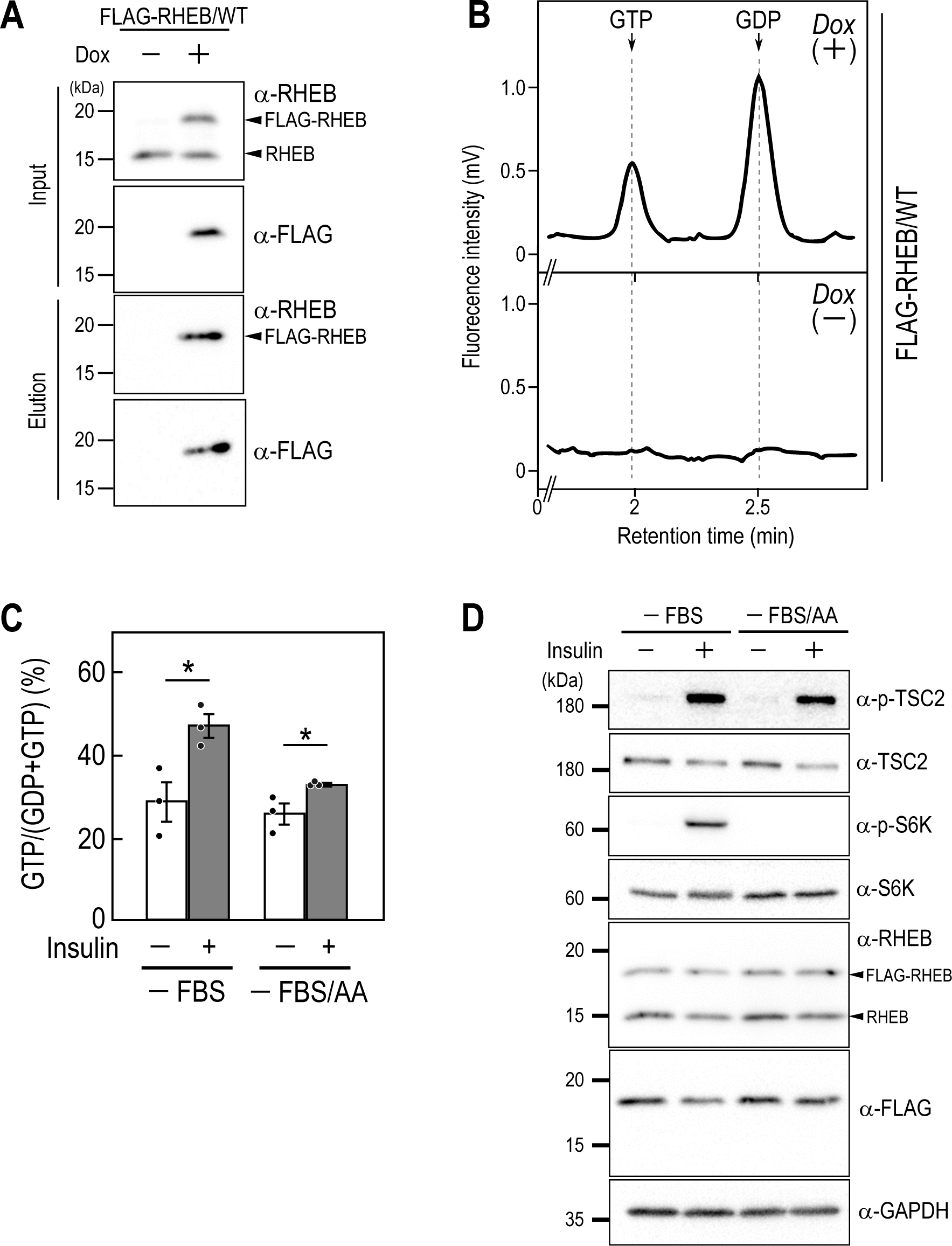
Fluor-HPLC Analysis of the activation states of FLAG-RHEB. (A) Dox-dependent expression of FLAG-tagged RHEB in HeLa cells. The cell line expressing FLAG-tagged RHEB proteins in a Dox-dependent manner was cultured in the absence or presence of 8 ng/ml of Dox for 24 h, and the cell lysates (*Input*) and immunoprecipitates (*Elution*) were subjected to western blot analysis using the indicated antibodies. (B) Representative chromatogram of guanine nucleotides bound to FLAG-tagged RHEB. Anti-FLAG immunoprecipitates from the indicated cells were subjected to Fluor-HPLC analysis. (*C, D*) Effect of insulin on the guanine nucleotide-bound states of FLAG-RHEB/WT and mTORC1 signaling. HeLa cells expressing FLAG-RHEB/WT were cultured for 24 h in FBS-depleted DMEM (*-FBS*) or for 3 h in Hank’s Balanced Salt Solution (*-FBS/AA*), followed by stimulation with 500 nM insulin for 10 min. Anti-FLAG immunoprecipitated from the cell lysates were subjected to Fluor-HPLC analysis, and the relative amounts of guanine nucleotides associated with FLAG-tagged proteins were quantified from the peak areas of GTP and GDP (*C*). Data represent the means ± SEM from three independent experiments and indicate individual data points; * *P* < 0.05 by Student’s *t*-test. Cell lysates prepared from cells treated under the indicated conditions were subjected to western blot analysis using the indicated antibodies (*D*).

### Assessing the Dynamics of RHEB Activity with Fluor-HPLC

The activation status of RHEB is tightly controlled, particularly in response to insulin stimulation, a process pivotal for cellular energy homeostasis (Menon et al., 2014). The tuberous sclerosis complex 2 (TSC2), which functions as a GTPase-activating protein (GAP) for RHEB, maintains RHEB in an inactive GDP-bound state under basal conditions. TSC2’s GAP activity is inactivated upon insulin stimulation through Akt-mediated phosphorylation, activating RHEB (Dibble et al., 2012; Inoki et al., 2003; Thomas and Hall, 1997). This active form of RHEB, bound to GTP, subsequently stimulates the mechanistic target of rapamycin complex 1 (mTORC1), a central regulator of protein synthesis and cell growth. mTORC1 activation is also regulated by amino acids, which are required to recruit mTORC1 to lysosomes, the compartments of mTORC1 activation. We investigated the activation dynamics of RHEB upon insulin stimulation in the presence or absence of amino acids by Fluor-HPLC (Fig. 2*C*). HeLa cells expressing FLAG-RHEB were subjected to serum starvation in the presence or absence of amino acids, followed by insulin stimulation, and the guanine nucleotides bound to FLAG-RHEB were analyzed by Fluor-HPLC. We found that insulin stimulation, particularly in the presence of amino acids (AA), led to an increase in the ratio of the GTP-bound form of FLAG-RHEB. This effect was also observed, though somewhat less, in the absence of amino acids. Western blot analysis revealed that TSC2 phosphorylation was induced upon insulin stimulation, but the extent of the phosphorylation was reduced in the absence of amino acids, consistent with previous studies (Fig. 2*D*) (Yang et al., 2020). Similarly, S6K (a substrate of mTORC1) was phosphorylated upon insulin stimulation in the presence of amino acids, but not in their absence. These findings show the utility of Fluor-HPLC in assessing alterations in RHEB activity following insulin stimulation.

### Evaluation of the Effect of KRAS/G12C Inhibitors on KRAS Activation Status by Fluor-HPLC

Next, we analyzed the small GTPase KRAS, mutations of which are frequently found in various malignancies. Among these, the KRAS/G12C mutation is particularly interesting due to its prevalence and impact on tumor development. Sotorasib, a novel small molecule inhibitor targeting KRAS/G12C, has emerged as a promising therapeutic option for KRAS G12C-positive non-small cell lung cancer (Canon et al., 2019). Mechanistically, Sotorasib binds specifically to the GDP-bound form of KRAS/G12C, thereby trapping KRAS/G12C in the inactive form and inhibiting KRAS/G12C-mediated oncogenic signaling. Studies assessing the efficacy of KRAS/G12C inhibitors, such as sotorasib, in cell cultures or mice have often involved monitoring ERK phosphorylation downstream of KRAS or using techniques like GST-pull-down assays. Nevertheless, the influence of sotorasib on the guanine-nucleotide binding status of KRAS/G12C within cellular contexts remains uncertain. To address this, we conducted Fluor-HPLC analysis on FLAG-KRAS expressed in HeLa cells (Fig. 3*A*). We found that the ratios of the GTP-bound form of FLAG-KRAS/WT and G12C were approximately 10% and 60%, respectively (Fig. 3*B*), consistent with previous observations suggesting KRAS/G12C as an active mutant. We then assessed the impact of sotorasib on the guanine nucleotide-bound form of KRAS (Fig. 3*C*). Sotorasib significantly decreased the ratio of the GTP-bound form of KRAS/G12C but not KRAS/wild-type and other cancer-associated KRAS mutants such as KRAS/G12D, G12V, and G13D. Consistent with the specific effect of sotorasib, it inhibited ERK phosphorylation induced by KRAS/G12C but not by other KRAS mutants (Fig. 3*D*). Additionally, the concentration-dependent inhibition of KRAS/G12C activity and ERK phosphorylation by sotorasib showed good correspondence (*SI Appendix*, Fig. S2 *A* and *B*) (Canon et al., 2019).

**Figure 3.**
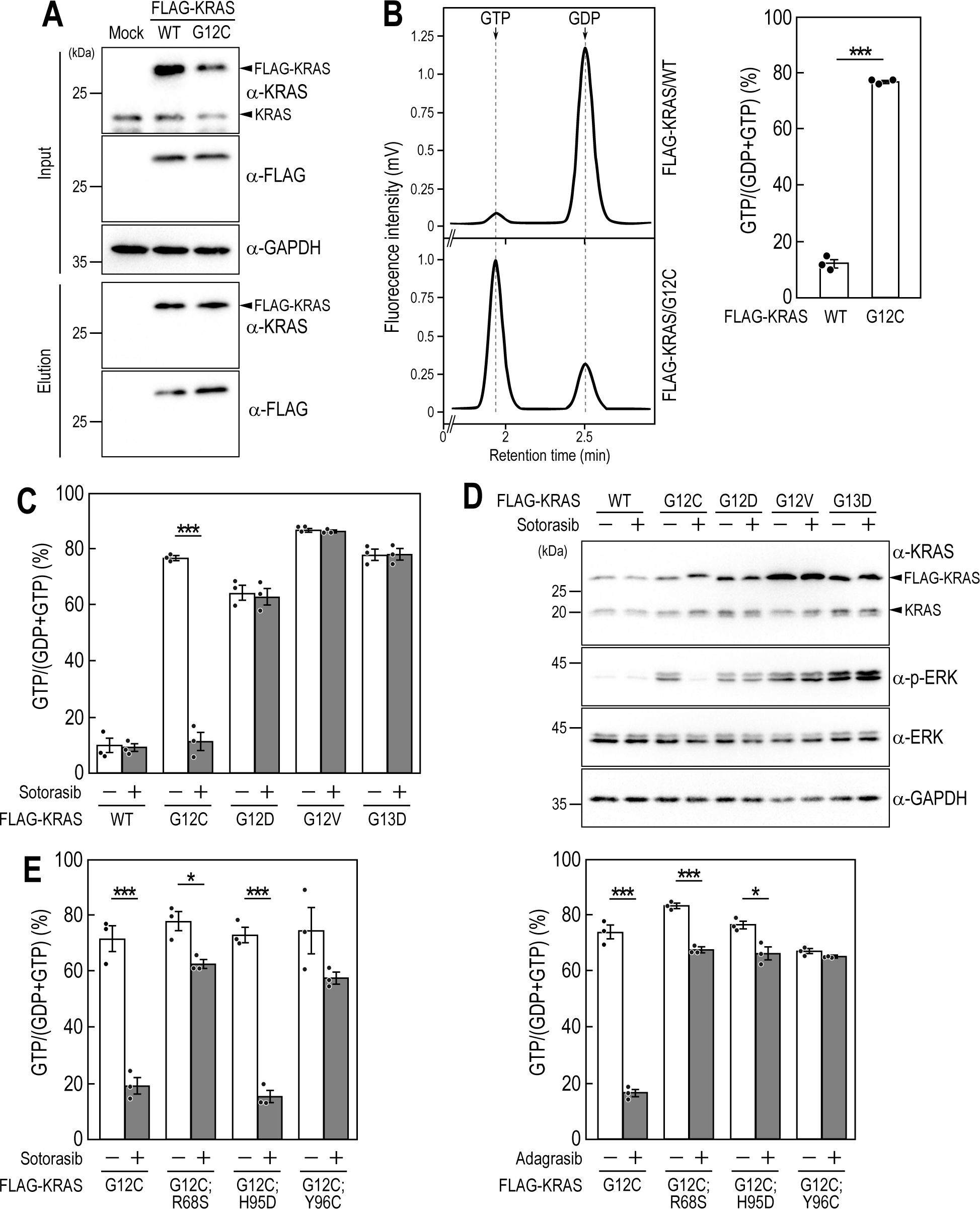
Analysis of the activation states of wild-type and mutants of FLAG-KRAS and the effects of KRAS/G12C inhibitors on their activation states. (*A, B*) Activation states of FLAG-KRAS wild-type and the G12C mutant. The total cell lysates (*Input*) and anti-FLAG immunoprecipitates (*Elution*) prepared from HeLa cells (*Mock*) and HeLa cells expressing FLAG-KRAS/WT or FLAG-KRAS/G12C in a Dox-dependent manner were subjected to western blot analysis using the indicated antibodies (*A*). Representative chromatogram of guanine nucleotides bound to FLAG-KRAS wild-type and the G12C mutant (*B, left panel*). The relative amounts of guanine nucleotides associated with FLAG-tagged proteins were quantified from the peak areas of GTP and GDP using Fluor-HPLC analysis (*B, right panel*). Data represent the means ± SEM from three independent experiments and indicate individual data points; *** *P* < 0.001 by Student’s *t*-test. (*C*) Effects of sotorasib on the activation states of wild-type and mutants of FLAG-KRAS. HeLa cells expressing wild-type or mutants of FLAG-KRAS were cultured for 24 h in the absence or presence of 1 µM sotorasib. Anti-FLAG immunoprecipitates from the cell lysates were subjected to Fluor-HPLC analysis, and the relative amounts of guanine nucleotides associated with FLAG-tagged proteins were quantified from the peak areas of GTP and GDP. Data represent the means ± SEM from three independent experiments and indicate individual data points; *** *P* < 0.001 by Student’s *t*-test. (*D*) Effects of sotorasib on the phosphorylation of ERK in HeLa cells expressing wild-type and mutants of FLAG-KRAS. Cell lysates prepared from the indicated cell lines were subjected to western blot analysis using the indicated antibodies. (*E*) Analysis of the effects of KRAS secondary mutations on the KRAS/G12C inhibitory activity of sotorasib and adagrasib. MIA-Paca2 cells expressing the indicated mutants of FLAG-KRAS were cultured for 24 h in the absence or presence of 1 µM sotorasib (*left panel*) or 1 µM adagrasib (*right panel*). Anti-FLAG immunoprecipitates from the cell lysates were subjected to Fluor-HPLC analysis, and the relative amounts of guanine nucleotides associated with FLAG-tagged proteins were quantified from the peak areas of GTP and GDP. Data represent the means ± SEM from three independent experiments and indicate individual data points; * *P* < 0.05; *** *P* < 0.001 by Student’s *t*-test.

Clinical studies on cancer therapy using KRAS/G12C inhibitors have revealed KRAS/G12C variants harboring secondary mutations conferring resistance to KRAS/G12C inhibitors (Awad et al., 2021; Tanaka et al., 2021). While both sotorasib and adagrasib are KRAS/G12C inhibitors, they have distinct chemical structures and binding modes to KRAS/G12C, leading to varying susceptibilities to secondary mutations in KRAS/G12C. For example, KRAS/G12C;R68S and KRAS/G12C;Y96C are resistant to both sotorasib and adagrasib, while KRAS/G12C;H95D is resistant to adagrasib but sensitive to sotorasib. We thus analyzed the effects of these mutations on KRAS activation status using MIA-Paca2, a pancreatic cancer-derived cell line. We found that the GTP-bound ratios of KRAS/G12C;R68S and KRAS/G12C;Y96C were minimally affected by either sotorasib or adagrasib, whereas sotorasib effectively suppressed the GTP-bound ratio of KRAS/G12C;H95D, which is consistent with the differential sensitivity observed in previous studies (Awad et al., 2021; Tanaka et al., 2021). Together, these findings show the utility of Fluor-HPLC analysis for assessing the guanine nucleotide-bound status of KRAS in cells and evaluating the efficacy of its inhibitors on its activity.

### Characterizing KRAS Activation States in Tumor Models Using Fluor-HPLC

Finally, we evaluated the utility of Fluor-HPLC for assessing KRAS activation status *in vivo* using a syngeneic metastatic B16F10 mouse melanoma model in C57/BL6 mice. FLAG-mEGFP-KRAS-expressing B16F10 melanoma cells were subcutaneously implanted into mice, and tumor tissue extracts were subjected to immunoprecipitation with FLAG antibody followed by Fluor-HPLC analysis of bound guanine nucleotides (Fig. 4*A*). Our findings revealed that approximately 20% of FLAG-mEGFP-KRAS/WT in tumor tissue was in the GTP-bound state. In comparison, approximately 60% of FLAG-mEGFP-KRAS/G12C and G12D exhibited GTP-bound status, consistent with results from cultured cells (Fig. 4*B*). Additionally, we investigated the impact of sotorasib administration on FLAG-mEGFP-KRAS activation in tumor tissues, demonstrating a reduction in the proportion of GTP-bound KRAS/G12C to approximately 20%. In contrast, the activation status of KRAS/G12D remained unaffected (Fig. 4*C*). These results underscore the potential of Fluor-HPLC in conjunction with *in vivo* models to discern KRAS activation dynamics and assess the efficacy of targeted therapeutics.

**Figure 4.**
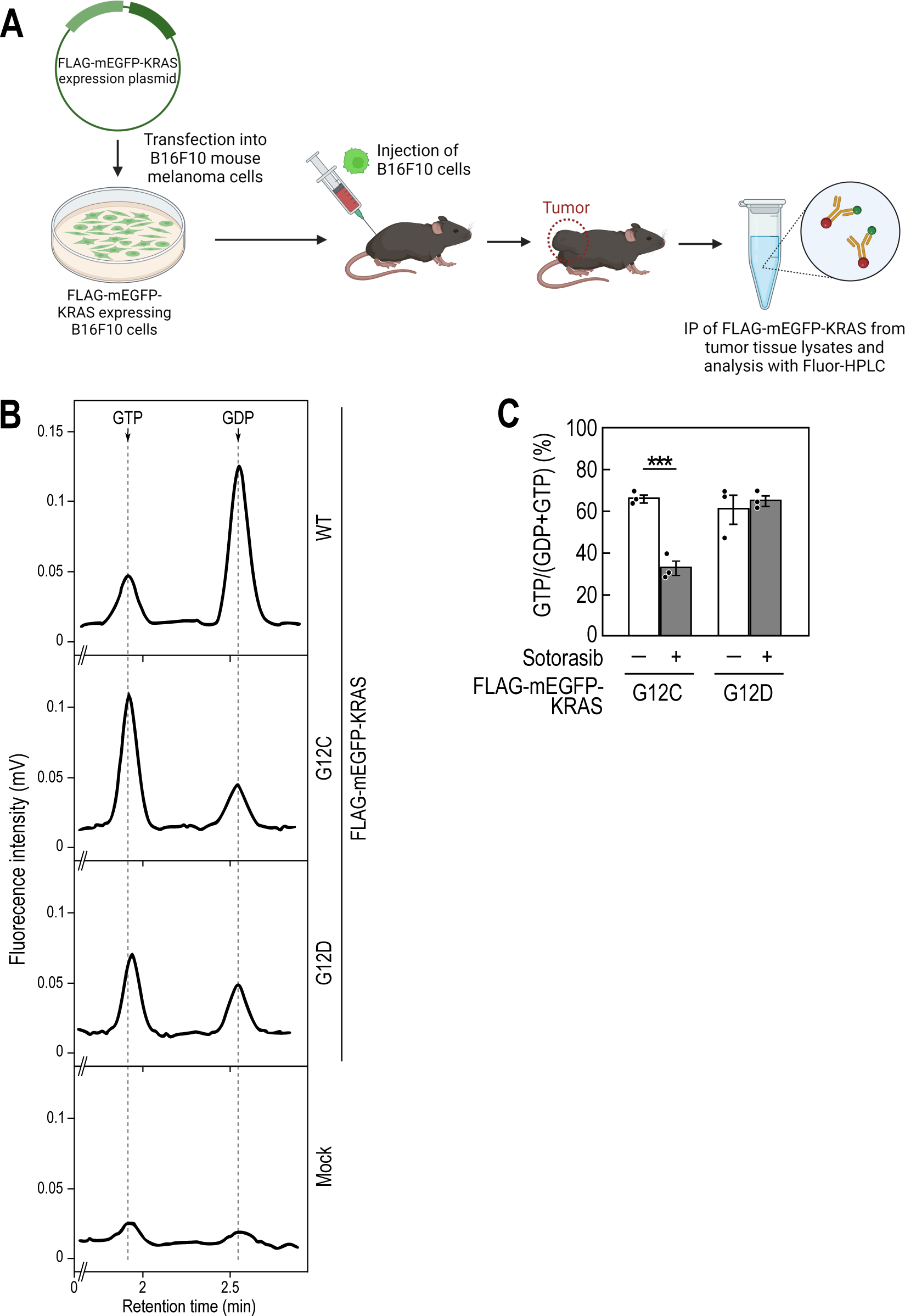
In vivo Analysis of FLAG-KRAS activation states with Fluor-HPLC. (*A*) Schematic diagram of the analysis of the activation state of FLAG-KRAS in tumor tissues using Fluor-HPLC in combination with a syngeneic mouse tumor model. Adapted from “Immunity Against Tumors is Specific”, by BioRender.com (2024). Retrieved from https://app.biorender.com/biorender-templates. (*B*) Representative chromatogram of guanine nucleotides bound to wild-type and mutants of FLAG-mEGFP-KRAS expressed in tumor tissues. Tumors derived from control B16F10 cells (*Mock*) or B16F10 cells expressing wild-type and mutants of FLAG-mEGFP-KRAS were harvested, and the tumor tissue extracts were subjected to anti-FLAG immunoprecipitation followed by Fluor-HPLC analysis. (*C*) Effect of sotorasib on the activation states of FLAG-KRAS mutants in tumor tissues. Mice with tumors expressing FLAG-KRAS/G12C or G12D were orally administered sotorasib (20 mg/kg) 2 h before harvesting tumor tissues. Anti-FLAG immunoprecipitates from the tumor tissue extracts were subjected to Fluor-HPLC analysis, and the relative amounts of guanine nucleotides associated with FLAG-tagged proteins were quantified from the peak areas of GTP and GDP. Data represent the means ± SEM from three independent experiments and indicate individual data points; *** *P* < 0.001 by Student’s *t*-test.

## Discussion

In this study, we established a highly sensitive and versatile fluorescence-detection HPLC method for assessing the activation states of small GTPases. This technique enables the analysis of guanine-nucleotide binding states even at endogenous expression levels of small GTPases. Using this approach, we investigated the activation state changes of RHEB in response to insulin stimulation and showed that the increase in its GTP-bound form is affected by amino acids in the culture medium. Additionally, we quantitatively assessed the effects of KRAS/G12C inhibitors on KRAS activation states in cultured cells and syngeneic mouse models.

Understanding the guanine-nucleotide binding state of small GTPases in cells is crucial for elucidating their functions and their associations with diseases due to genetic mutations. HPLC has been previously employed to analyze guanine nucleotide binding to small GTPases (Ahmadian et al., 1999; Araki et al., 2021; Araki and Kontani, 2024; Bromberg et al., 1994; Feuerstein et al., 1987; Hannan et al., 2021; Hanzal-Bayer et al., 2002; John et al., 1990; Lawrence et al., 2019; Scrima et al., 2008; Sot et al., 2013; Su et al., 2020; Weiss et al., 1989). However, since these studies relied on UV detection of guanine nucleotides, relatively large quantities of small GTPases (e.g., purified recombinant proteins) were required to detect guanine nucleotides effectively. Our study, by employing fluorescence derivatization of guanine nucleotides, achieved over 100-fold higher sensitivity compared to UV detection, enabling quantitative analysis of guanine nucleotides bound to small GTPases even at their endogenous expression levels (Fig. 2 and 3). To our knowledge, only about 10% of the ∼150 small GTPases in humans have been analyzed for their activation states at endogenous levels in the cell (Rooij and Bos, 1997; Sander et al., 1998; Santy and Casanova, 2001). Thus, the Fluor-HPLC method will provide a valuable tool for analyzing the activation states and their regulation of various small GTPases that have yet to be characterized. Furthermore, this technique can detect subtle changes in guanine-nucleotide binding states, offering superior quantitative and reproducible analysis compared to western blotting-based quantitative analysis such as GST pull-down assays.

Activated (GTP-bound) RHEB triggers the activation of mTORC1 at the lysosome, which in turn phosphorylates various proteins, including S6K (Inoki et al., 2003; Long et al., 2005; Saucedo et al., 2003; Stocker et al., 2003). Our data showed that, despite 30% of intracellular RHEB being in the GTP-bound form, S6K phosphorylation was not detected without insulin stimulation (Fig. 2 *C* and *D*). Given the predominant cytosolic localization of RHEB (Angarola and Ferguson, 2019), this lack of S6K phosphorylation likely occurred because most of the GTP-bound RHEB existed in the cytosol and did not significantly contribute to mTORC1 activation at the lysosome membrane (44). Upon insulin stimulation, TSC2, a RHEB GTPase-activating protein, dissociates from the lysosome membrane (Menon et al., 2014), which should increase the amount of membrane-localized GTP-bound RHEB. Considering that the proportion of membrane-localized RHEB is very low relative to the total amount of RHEB in the cell, it is not surprising that the extent of the increase in the proportion of GTP-bound RHEB upon insulin stimulation, when considering the entire cell, was smaller compared to the marked increase in phosphorylated S6K (23). Future analysis enabling the measurement of guanine-nucleotide binding states of membrane-localized RHEB alone could provide detailed insights into RHEB activation dynamics upon insulin stimulation.

The degree of RHEB activation in response to insulin varied with amino acids in the medium: a 1.7-fold increase under serum starvation alone versus a 1.3-fold increase under both serum and amino acid starvation (Fig. 2*C*). This difference might be due to variations in TSC2 localization to the lysosome membrane. Previous studies suggest that TSC2 localizes more to the lysosome membrane during amino acid starvation due to a reduction in the degree of its phosphorylation (Yang et al., 2020). Indeed, our study showed lower TSC2 phosphorylation under serum and amino acid starvation compared to serum starvation alone during insulin stimulation (Fig. 2*D*). Thus, more TSC2 localizing to the lysosome under amino acid starvation could efficiently inactivate RHEB, reducing its activation upon insulin stimulation. Furthermore, it has been reported that amino acid starvation lowers intracellular calcium ion levels, alleviating the inhibition of TSC2 by calmodulin and enhancing TSC2’s ability to act on RHEB (Amemiya et al., 2021). Thus, calmodulin-regulated TSC2 activity might also play a role in the inactivation of RHEB during amino acid starvation.

Previous evaluations of sotorasib efficacy in cultured cells used pull-down assays to measure GTP-bound KRAS levels and ERK phosphorylation (Canon et al., 2019; Ryan et al., 2022). Here, we demonstrated that sotorasib decreases GTP-bound KRAS/G12C while increasing GDP-bound KRAS/G12C in cells, aligning with in vitro studies showing that sotorasib acts on GDP-bound KRAS/G12C to inhibit guanine nucleotide exchange. Simultaneously measuring both the GTP-bound and GDP-bound forms of small GTPases will provide more valuable insights into the regulatory mechanisms of their activation states and the action mechanisms of compounds targeting small GTPases.

We have also shown that Fluor-HPLC assay can be used to evaluate sotorasib activity against KRAS *in vivo* (Fig. 4). This method is also applicable to xenograft mouse models commonly used in anti-tumor efficacy studies of sotorasib (Canon et al., 2019). Combining these mouse models with Fluor-HPLC can deepen our understanding of the correlation between KRAS activation states and tumor formation. Additionally, provided that immunoprecipitation is possible, this method could be applied to other small GTPases to analyze the effects of stress and disease conditions on their activation states in various tissues at the whole-organism level.

While small GTPases generally exist in GDP- or GTP-bound states in cells, some studies indicate that particular small GTPases may exist in nucleotide-free forms within cells. Small GTPases are generally considered unstable and often inactivate in nucleotide-free forms, yet ARF1 and RAB8 have been shown to maintain stability and activity by interacting with phosphatidylinositol 4,5-bisphosphate and the chaperone Mss4, respectively (Itzen et al., 2006; Nuoffer et al., 1997; Terui et al., 1994). Furthermore, specific monobodies targeting KRAS can bind to nucleotide-free KRAS in cells, thereby inhibiting the guanine-nucleotide exchange and KRAS-mediated signaling (Khan et al., 2022). Nevertheless, these studies did not directly quantify the nucleotide-free forms of small GTPases they examined. By combining absolute quantification of immunoprecipitated proteins with Fluor-HPLC, we could achieve a stoichiometric evaluation of guanine-nucleotide binding states of small GTPases, allowing for quantitative analysis of nucleotide-free forms.

In summary, this study introduces a robust fluorescence-detection HPLC technique for precisely quantifying guanine nucleotide states in small GTPases. By offering detailed quantification of guanine nucleotide-binding states, this method provides a clearer understanding of the activation and inactivation mechanisms of small GTPases, thus enhancing our capacity to study small GTPase functions and assess the impact of therapeutic agents.

## Materials and Methods

### Reagents and Solutions

HPLC-grade acetonitrile and HPLC-grade tetrahydrofuran were purchased from Kanto Chemical. Phosphatase Inhibitor Cocktail (EDTA free) and HPLC-grade SDS were purchased from Nacalai tesque. Sotorasib, adagrasib, and PEG300 were purchased from Selleck Chemicals. Restriction enzymes were purchased from New England Biolabs. 3,4-Dimethoxyphenylglyoxal hydrate (DMPG, Alfa Aesar) was dissolved in DMSO-water mixture (2:3, v/v). GDP and GTP (Sigma-Aldrich) were dissolved in water, with concentrations determined using their molecular absorption coefficients. Ganciclovir and all other chemicals were purchased from Fujifilm Wako Pure Chemicals.

### Reversed-Phase HPLC Analysis with Fluorescence Detection (Fluor-HPLC Analysis)

The Fluor-HPLC system comprised a DGU-20A3 degasser, an LC-20AD pump, a SIL-20AC autosampler, a CTO-20AC column oven, an RF-20A fluorescence detector, and a CBM-20A Communication Bus Module (Shimadzu). Instrumental control and data analysis were performed using LC Solution (Shimadzu). Fluorescent derivative guanine nucleotides were separated using a Gemini 3 µm NX-C18 110 Å LC Column (100 × 3 mm) (Phenomenex, 00D-4453-Y0) with a Gemini NX C18 Security Guard Cartridge (4.0 × 3.0 mm) (Phenomenex, AJ0-8368). Mobile phase A consisted of 2.8% acetonitrile, 1.4% tetrahydrofuran, and 12.5 mM potassium phosphate buffer (pH 8.0). Mobile phase B was 100% acetonitrile. The gradient program was as follows: 0% B (0 min)−0% B (1.8 min)−10% B (1.85 min)−10% B (2 min)−12% B (2.05 min)−12% B (4.4 min)−0% B (4.45 min)−0% B (9 min). Elution was performed at room temperature with a flow rate of 1.0 ml/min. The detection wavelength was set at an excitation wavelength of λex = 400 nm and an emission wavelength of λem = 510 nm.

### Plasmids

Plasmids for establishing stable cell lines were constructed using the Gateway cloning technology (Thermo Fisher Scientific). pENTR-TiTRE and pENTR-TRE3G were used as donor plasmids for tightly controlled gene expression by tetracycline-responsive promoters. A donor plasmid for the expression of FLAG-tagged RHEB was constructed as described previously (Araki et al., 2021). Donor plasmids for the expression of wild-type and mutants of FLAG-KRAS were constructed by cloning each gene into the EcoRI/SalI site of pENTR-TRE3G vector. Expression Plasmids were obtained by LR clonase (Thermo Fisher Scientific) reaction of each donor plasmid with the destination plasmid pAAVS1_Puro_ccdb_Ubc_rtTA (Araki et al., 2021), according to the manufacturer’s protocol. Plasmids for expression of wild type and mutants of FLAG-mEGFP-KRAS were constructed by cloning each gene into the HindIII/BamHI site of R26-N-FLAG-mEGFP vector, which was constructed by replacing the EGFP-coding region of R26-EGFP HMEJ donor vector with a coding DNA fragment of FLAG-mEGFP. R26-EGFP HMEJ donor vector was a gift from Hiroshi Kiyonari (Addgene plasmid # 137927; http://n2t.net/addgene:137927; RRID:Addgene_137927). The pCas-Guide-AAVS1-T2 and pCas-Guide-ROSA26-1 plasmids were constructed by cloning AAVS1-T2 and ROSA26-1 gRNA fragment, respectively, into the pCas-Guide vector (OriGene). The target sequences of gRNAs are provided in *SI Appendix*, Table S1.

### Western Blotting

Western blotting was performed as described previously (Araki et al., 2021). Cell lysates were separated on SDS-polyacrylamide gels and transferred to ClearTrans SP PVDF Membranes (Fujifilm Wako Pure Chemical) using a Trans-Blot Turbo Transfer System (BIORAD). After blocking with 3% skim milk or 5% BSA (for anti-phospho-S6K and anti-phospho-ERK1/2) in TBS containing 0.1% Tween 20, membranes were probed with antibodies and subjected to chemiluminescent measurement using EzWest LumiOne (ATTO) and LuminoGraph II (ATTO). Antibodies are listed in *SI Appendix*, Table S2.

### Cell Culture and Establishment of Stable Cell Lines

HeLa, MIA-Paca2, and B16F10 cells were cultured in Dulbecco’s Modified Eagle’s Medium (DMEM) containing 10% FBS, penicillin, and streptomycin. Isogenic stable cell lines of HeLa and MIA-Paca2 expressing genes of interest were generated by CRISPR/Cas9-driven targeted integration of the gene into the safe-harbor genomic locus *AAVS1*, as described previously (Araki et al., 2021; Dalvai et al., 2015). B16F10 cells expressing genes of interest were generated by CRISPR/Cas9-driven targeted integration of the gene into the *ROSA26* locus, as described previously (Abe et al., 2020). Briefly, B16F10 cells were co-transfected with expression plasmid and pCas-Guide-ROSA26-1 plasmid. Forty-eight hours post-transfection, cells were harvested and GFP-positive cells were isolated using the Cell Sorter MA900. After 2 to 3 passages, the cells were re-sorted to further enrich the GFP-positive population.

### Immunoprecipitation of Small GTPases for Fluor-HPLC Analysis

Anti-DYKDDDDK (anti-FLAG) tag antibody-conjugated Dynabeads protein G (Thermo Fisher Scientific) were prepared as follows: Dynabeads protein G (20 μl) was washed with 250 μl of PBS containing 0.1% BSA and resuspended in 50 μl of PBS containing 0.1% BSA. The washed beads were incubated with 2 µg of anti-DYKDDDK tag antibody at room temperature for 30 min with gentle mixing. After washing with 250 μl of ice-cold wash buffer-1 (40 mM Tris-HCl, pH 7.5, 100 mM NaCl, 5 mM MgCl_2_, 0.1% (w/v) Triton X-100, and 1 mM DTT), beads were suspended in 50 μl of ice-cold wash buffer-1 and used for the subsequent immunoprecipitation experiments. Cells cultured in a 100-mm or 150-mm dish were lysed with 550 μl of ice-cold extraction buffer (40 mM Tris-HCl, pH 7.5, 100 mM NaCl, 5 mM MgCl_2_, 1% (w/v) Triton X-100, 1 mM DTT, 0.2% phosphatase inhibitor cocktail (EDTA free), and 0.5 mM AEBSF). Lysates were centrifuged at 15,000 rpm at 4°C for 5 min, and the supernatants were incubated with anti-DYKDDDK tag antibody-conjugated beads at 4°C for 30 min. Beads were washed twice with 400 μl of ice-cold wash buffer-1 and suspended with 250 μl of ice-cold wash buffer-2 (30 mM sodium phosphate buffer, pH 7.0, 100 mM NaCl, 1 mM MgCl_2_, and 0.1% (w/v) Lubrol-PX) and transferred to a Protein LoBind tube (Eppendorf). After removing the buffer, beads were suspended with 70 μl of elution buffer (10 mM sodium phosphate buffer, pH 7.0, 0.1% SDS (HPLC-grade), 50 nM ganciclovir) and incubated at 90°C for 3 min. The supernatants were filtered using a Nanosep 3K Omega device (Pall), and the flow-through fraction was collected for fluorescence derivatization reaction of guanine nucleotides and ganciclovir as follows: sample (18 μl) was mixed with 100 mM DMPG (6 μl) and incubated at 37°C for 5 min under shaded light. The reactants were centrifuged at 15,000 rpm at 4°C for 1 min, and supernatants (10 μl) were subjected to Fluor-HPLC analysis.

### Insulin Stimulation of HeLa Cells

HeLa cells expressing FLAG-RHEB were starved for 24 h in FBS-depleted DMEM (DMEM supplemented with 10 mM HEPES, pH7.4, 0.16% sodium bicarbonate, 0.58 mg/ml glutamine and 0.1% fatty acid-free BSA) or for 3 h in Hank’s Balanced Salt Solution (Shimadzu Diagnostics). After stimulation of insulin (500 nM) for 10 min, cells were lysed as described above, and the cell lysates and anti-FLAG immunoprecipitates from the lysates were subjected to western blotting analysis and Fluor-HPLC analysis, respectively.

### Animal Studies

All animal experiments were performed according to protocols approved by the Meiji Pharmaceutical University Laboratory Animal Ethics Committee. Six-week-old female C57BL/6JmsSlc mice (Japan SLC) were housed in an environmentally-controlled room on a 12-hour light/dark cycle.

### Syngeneic Mouse Tumor Model

Cell suspensions of B16F10 cells or B16F10 cells expressing wild-type or mutants of FLAG-mEGFP-KRAS [4 × 10^6 cells/ml in PBS (−)] were injected subcutaneously into the flanks of mice.

At 10 days post-implantation, tumor tissues (∼200 mm^3^, calculated using the following formula: length × width^2^ × 0.52) were harvested and homogenized with 500 μl of ice-cold extraction buffer (40 mM Tris-HCl, pH 7.5, 100 mM NaCl, 5 mM MgCl_2_, 1% (w/v) Triton X-100, 1 mM DTT, 0.2% phosphatase inhibitor cocktail (EDTA free), and 0.5 mM AEBSF) using a BioMasher II homogenizer and vortexed at 4°C for 5 min. The tissue homogenates were centrifuged at 15,000 rpm at 4°C for 5 min, and the supernatants were subjected to anti-FLAG immunoprecipitation followed by Fluor-HPLC analysis. Sotorasib (dissolved in 5% DMSO, 40% PEG300, and 5% Tween 80) was administered orally (20 mg/kg) 2 h before harvest of tumor tissues.

## Acknowledgments

This work was supported in part by the Japan Society for the Promotion of Science (JSPS) KAKENHI (grant numbers 20K06565 and 23K05685 to KK).

## Author Contributions

M.A., T.K., and K.K. designed research; M.A., Y.K., and K.Y. performed research; M.A., Y.K., K.Y. and K.K. analyzed and interpreted data; and M.A. T.K., and K.K. wrote the paper.

## Notes

### Competing Interest Statement

The authors have declared no competing interest.

