## Supplementary Figures and Tables for "Quantifying small GTPase activation status using a novel fluorescence HPLC-based assay"

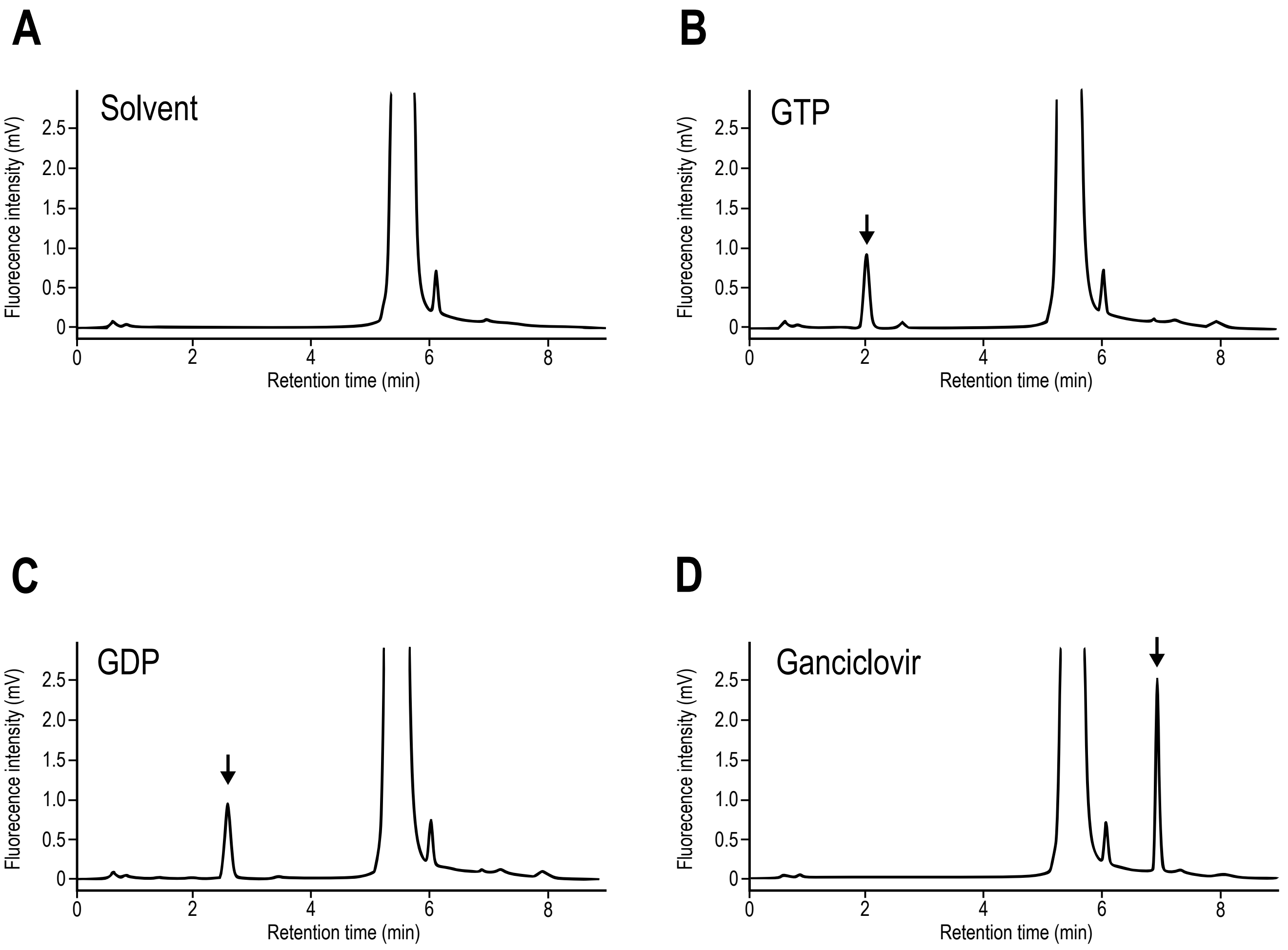


**Figure S1. Fluor-HPLC chromatograms of GTP, GDP, and ganciclovir.**
Representative chromatograms of solvent (*A*), 200 fmol GTP (*B*), 200 fmol GDP (*C*), and 375 fmol ganciclovir (*D*) obtained by Fluor-HPLC. The compounds were detected using a fluorescence detector at an excitation wavelength of 400 nm and an emission wavelength of 510 nm.

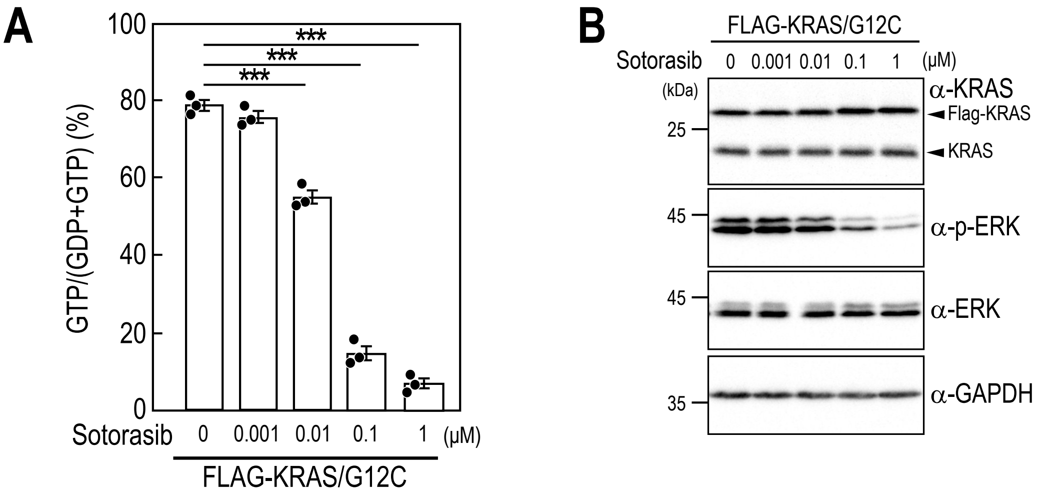


**Figure S2. Concentration-dependent impact of sotorasib on FLAG-KRAS/G12C activation states and ERK phosphorylation.**

(*A*) Effects of sotorasib on the activation states of FLAG-KRAS/G12C. HeLa cells expressing FLAG-KRAS/G12C were treated with the indicated concentration of sotorasib for 24 h, and anti-FLAG immunoprecipitates from the cell lysates were subjected to Fluor-HPLC analysis. The relative amounts of guanine nucleotides associated with FLAG-tagged proteins were quantified from the peak areas of GTP and GDP. Data represent the means ± SEM from three independent experiments and indicate individual data points; *** $P < 0.001$ by Dunnett's test. (*B*) Effects of sotorasib on ERK phosphorylation in HeLa cells expressing FLAG-KRAS/G12C. Cell lysates prepared from cells treated with the indicated concentration of sotorasib were subjected to western blot analysis using the indicated antibodies.

**Table S1. gRNAs used in this study**

| Name | Sequence | Reference |
| --- | --- | --- |
| AAVS1-T2 | 5'- GGGGCCACTAGGGACAGGAT -3' | P. Mali, et al., RNA-Guided Human Genome Engineering via Cas9. *Science* **339**, 823–826 (2013). |
| ROSA26-1 | 5'- ACTCCAGTCTTTCTAGAAGA -3' | V. T. Chu, et al., Increasing the efficiency of homology-directed repair for CRISPR-Cas9-induced precise gene editing in mammalian cells. *Nat. Biotechnol.* **33**, 543–548 (2015). |

**Table S2. Antibodies used in this study**

| Antibodies [ Working dilution ] | Source | Identifier |
|---|---|---|
| RHEB [ 1:2000 ] | Cell Signaling Technology | #13879, RRID:AB_2721022 |
| S6K [ 1:2000 ] | Cell Signaling Technology | #2708, RRID:AB_390722 |
| Phospho-S6K (Thr389) [ 1:2000 ] | Cell Signaling Technology | #9234, RRID:AB_2269803 |
| TSC2 [ 1:2000 ] | Cell Signaling Technology | #4308, RRID:AB_10547134 |
| Phospho-TSC2 (Thr1462) [ 1:2000 ] | Cell Signaling Technology | #3617, RRID:AB_490956 |
| ERK1/2 [ 1:2000 ] | Cell Signaling Technology | #4695, RRID:AB_390779 |
| Phospho-ERK1/2 (Thr202/Thr204) [ 1:2000 ] | Cell Signaling Technology | #4370, RRID:AB_2315112 |
| DYKDDDDK (FLAG) [ 1:2500 ] | Fujifilm Wako Pure Chemical | 014-22383, RRID:AB_10659717 |
| GAPDH [ 1:3000 ] | Fujifilm Wako Pure Chemical | 016-25523, RRID:AB_2814991 |
| HRP-conjugated anti-rabbit IgG [ 1:20000 ] | Jackson ImmunoResearch Labs | 111-035-144, RRID:AB_2307391 |
| HRP-conjugated anti-mouse IgG [ 1:20000 ] | Jackson ImmunoResearch Labs | 115-035-146, RRID:AB_2307392 |
